## Supplementary material for "High-level expression of the monomeric SARS-CoV-2 S protein RBD 320-537 in stably transfected CHO cells by the *EEF1A1*-based plasmid vector": Raw image data

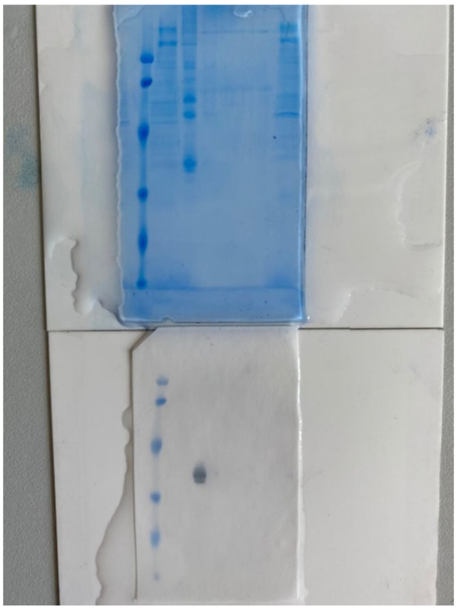

Fig_1B_Fig_1C - WhatsApp Image 2020-06-12 at 15.14.24.jpeg

11 june 2020
RBDv1 SDS-PAGE 12.5% and blot, purification on iminodiacetic acid (IDA)
lane 1: Prestained Protein Molecular Weight Marker #26612, 5 uL
lane 2: non-binding fraction, 5 uL concentrate
lane 3: 50 mM imidazole elution, 5 uL concentrate
lane 4: 50 mM imidazole elution (tail), 5 uL concentrate
lane 5: 100 mM imidazole elution, 5 uL concentrate
lane 6: 250 mM imidazole elution, 5 uL concentrate
lane 7: Na-EDTA elution, 5 uL concentrate
lane 8: harvested medium, 5 uL concentrate

All samples are 20x concentrates through Vivaspin PES membrane 10000 MWCO (Sartorius, UK, VS0102), SDS-PAGE 12.5% in reducing conditions (50 mM DTT)

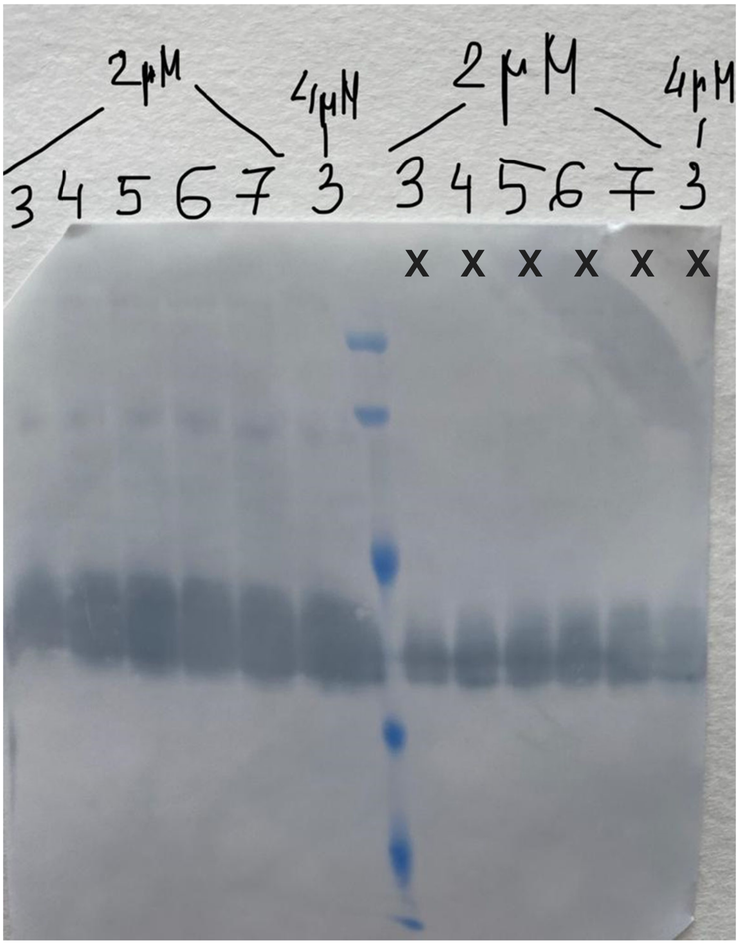


Fig_1D - d585c824-6486-486b-87c3-39b542531e7b.jpg

23 july 2020
RBDv1 SDS-PAGE 12.5%
lanes 1-6: 2 uM RBDv1 culture, 10 uL concentrate from days 3 to 8 (batch process)
lane 7: Prestained Protein Molecular Weight Marker #26612 5 uL
lanes 8-12: 2 uM RBDv1 culture, 1 uL concentrate from days 3 to 7 (batch process) – non-relevant data
lane 13: 4 uM RBDv1 culture, 1 uL concentrate from day 3 (batch process) – non-relevant data

All samples are 20x concentrates through Vivaspin PES membrane 10000 MWCO (Sartorius, UK, VS0102), SDS-PAGE 12.5% in reducing conditions (50 mM DTT)

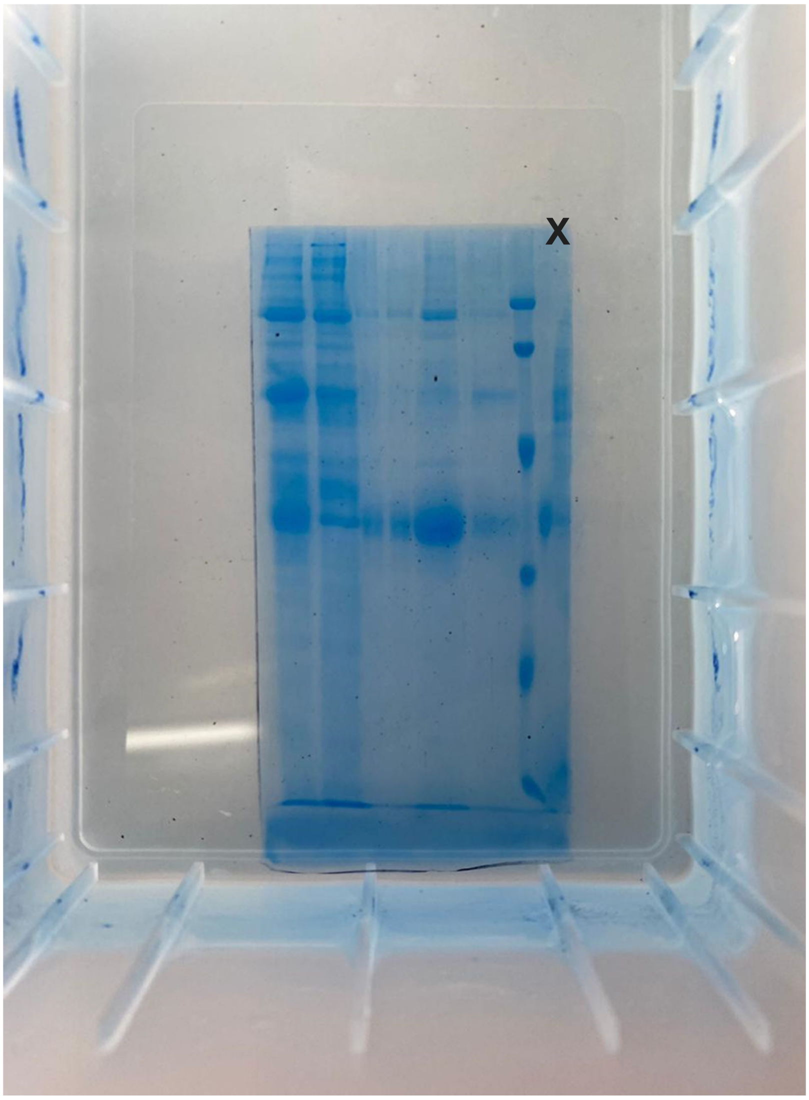


Fig _1E. e4c3a97b-5a15-442f-a852-0a600ea1cc47.jpg

17 august 2020
RBDv1 SDS-PAGE 12.5%, purification on nitrilotriacetic acid (NTA)
lane 1: harvested medium, 5 uL concentrate
lane 2: non-binding fraction, 5 uL concentrate
lane 3: 50 mM imidazole elution, 5 uL concentrate
lane 4: 50 mM imidazole elution (tail), 5 uL concentrate
lane 5: 250 mM imidazole elution, 5 uL concentrate
lane 6: Na-EDTA elution, 5 uL concentrate
lane 7: Prestained Protein Molecular Weight Marker #26612, 5 uL
lane 8: 250 mM imidazole elution, 0.5 uL concentrate – non-relevant data

All samples are 20x concentrates through Vivaspin PES membrane 10000 MWCO (Sartorius, UK, VS0102), SDS-PAGE 12.5% in reducing conditions (50 mM DTT)

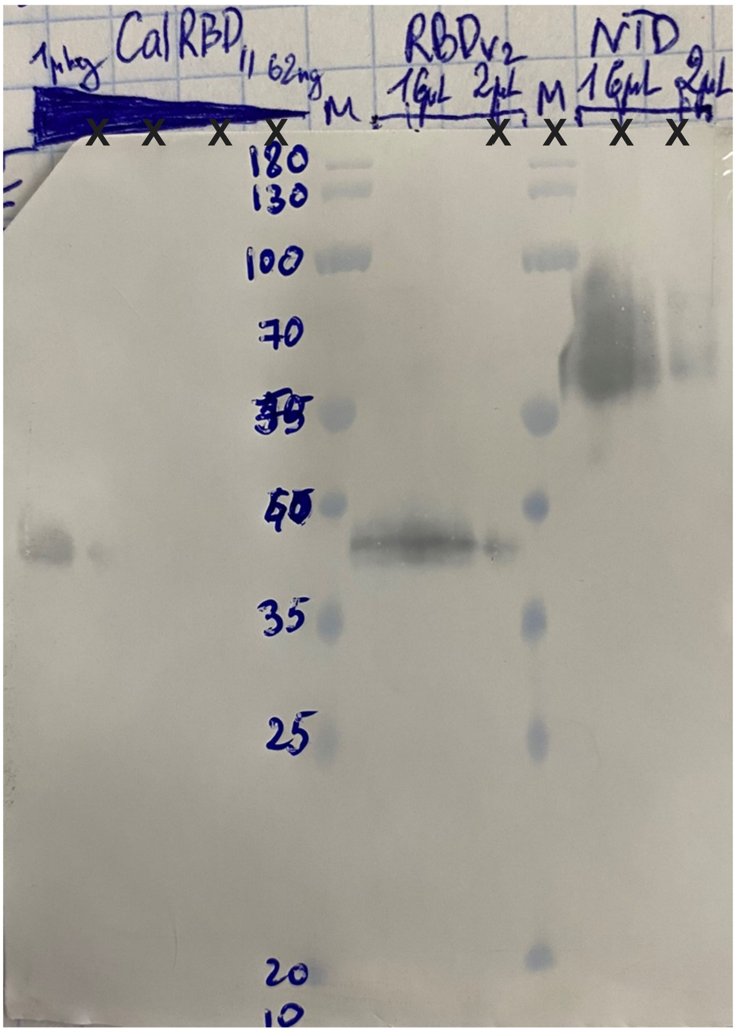


Fig 2_B - CoV_WB_20200908_RBDv2_NTD_2uM.jpg

17 august 2020
RBDv1 blot
lanes 1-5: RBDv1 purified calibrator, batch 11, 1 ug to 62 ng per well
lane 6: PageRuler Prestained Protein Ladder #26616, 3 uL
lane 7: 2 uM RBDv2 culture, 10 uL concentrate
lane 8: 2 uM RBDv2 culture, 1 uL concentrate – non-relevant data
lane 9: PageRuler Prestained Protein Ladder #26616, 3 uL
lane 10: 2 uM NTD culture, 10 uL concentrate – non-relevant data
lane 11: 2 uM NTD culture, 1 uL concentrate – non-relevant data

RBDv2 and NTD samples are 20x concentrates through Vivaspin PES membrane 10000 MWCO (Sartorius, UK, VS0102), SDS-PAGE 12.5% in reducing conditions (50 mM DTT)

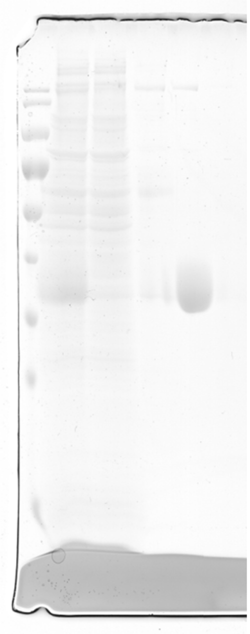

Fig 2_C - CoV_SDS_PAGE_20201002_RBDv2_PR03.tif

28 september  2020
RBDv2 SDS-PAGE 12.5%, 1L flask, 2 uM culture, purification on nitrilotriacetic acid (NTA), batch 03
lane 1: PageRuler Prestained Protein Ladder #26616, 5 uL
lane 2: harvested medium, 10 uL
lane 3: non-binding fraction, 10 uL
lane 4: 50 mM imidazole, 4 uL
lane 5: 250 mM imidazole elution, 2 uL
lane 6: Na-EDTA elution, 4 uL

Samples were loaded without concentration, the loaded volumes were proportional to the volumes of unconcentrated fractions. SDS-PAGE 12.5% in reducing conditions (50 mM DTT)

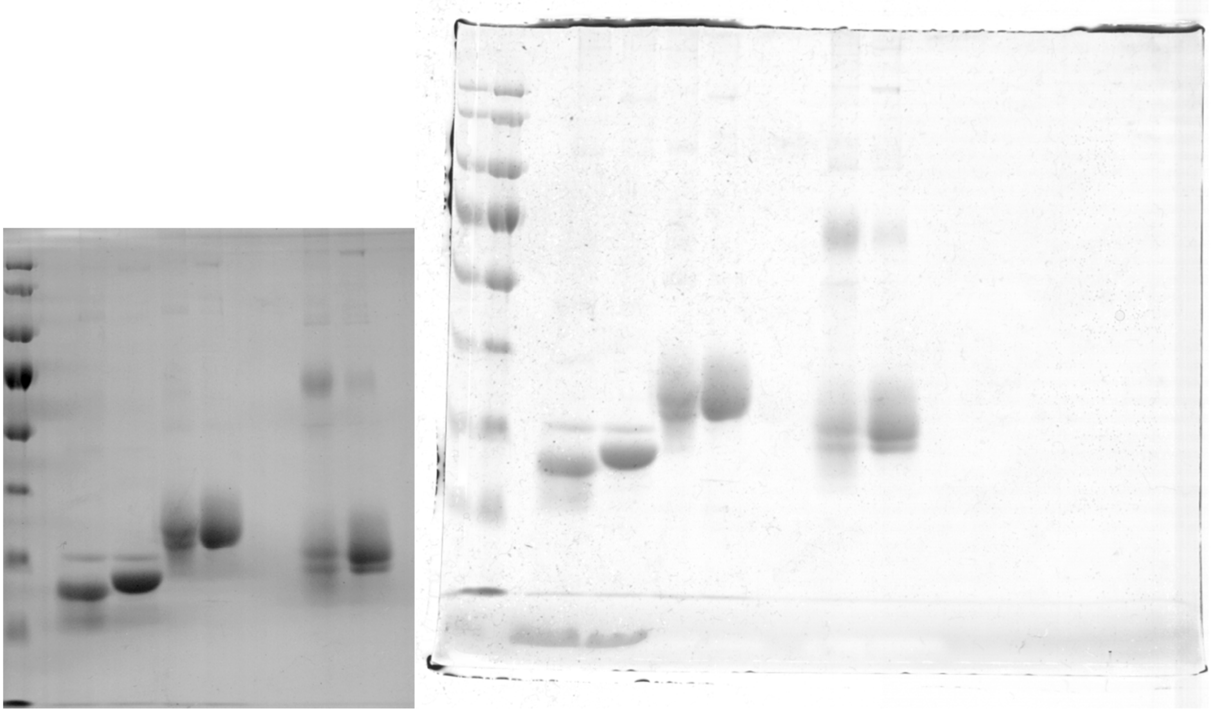

Fig 4_A - CoV_SDS_PAGE_20201014_RBD.tif

14 october 2020
RBDv1 vs RBDv2 SDS-PAGE 12.5%, PNGase treatment
lane 1: PageRuler Prestained Protein Ladder #26616, 5 uL
lane 2: X
lane 3: RBDv1, +PNGase F, +DTT , 5 ug
lane 4: RBDv2, +PNGase F, +DTT , 5 ug
lane 5: RBDv1, -PNGase F, +DTT , 5 ug
lane 6: RBDv2, -PNGase F, +DTT , 5 ug
lane 7: X
lane 8: X
lane 9: RBDv1, -PNGase F, -DTT , 5 ug
lane 10: RBDv2, -PNGase F, -DTT , 5 ug

For all samples volumes were adjusted to the final volume of 15 uL with 4x DualColor loading buffer (Fermentas, #R1011)
