## Supporting figures and tables for "High-level expression of the monomeric SARS-CoV-2 S protein RBD 320-537 in stably transfected CHO cells by the *EEF1A1*-based plasmid vector"

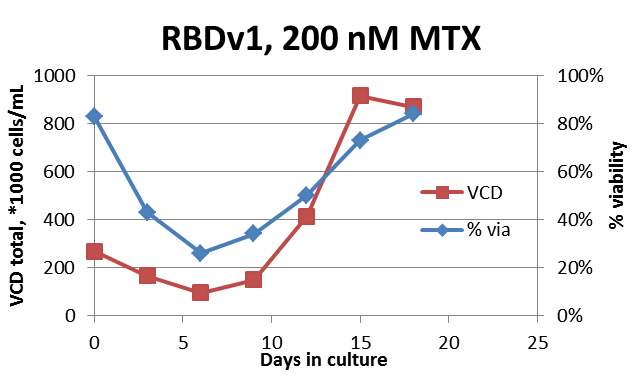

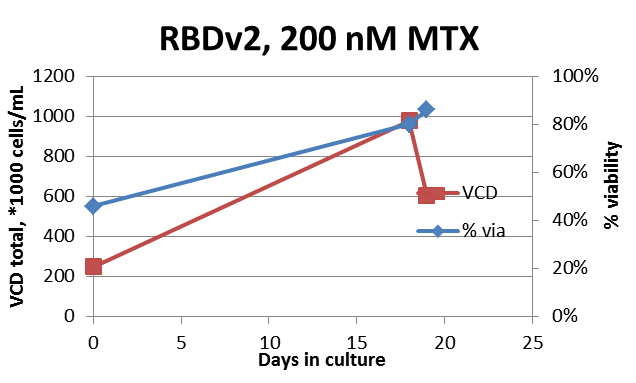

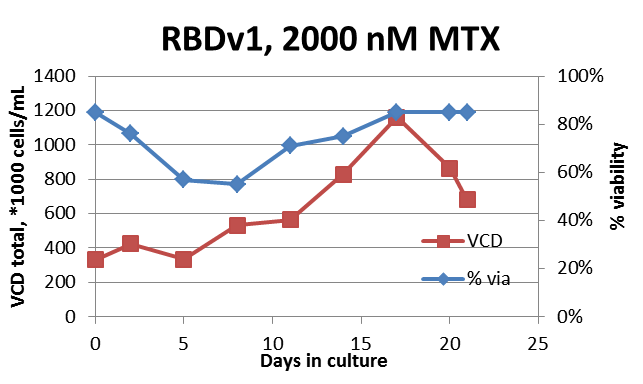

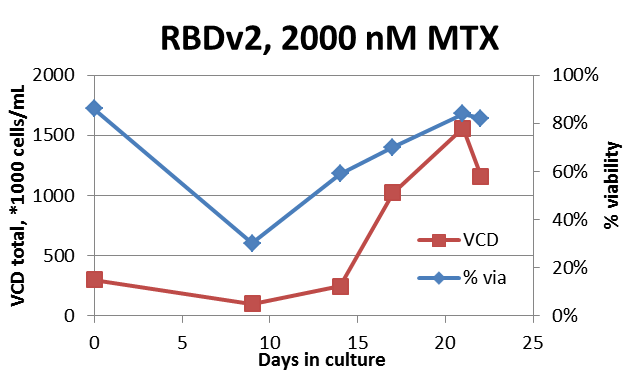

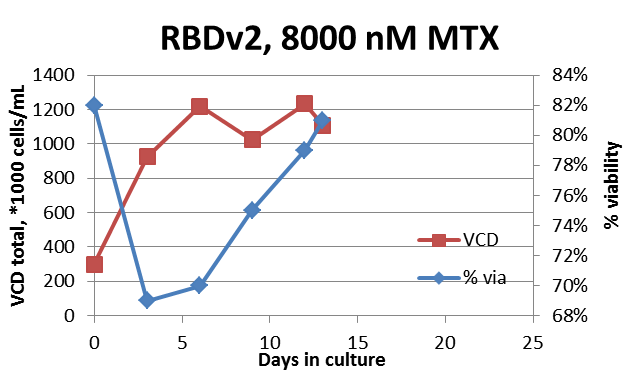

Supporting Figure S1. Cell growth and viability dynamics of initial selection and MTX-driven target gene amplification. RBDv1 – plasmid p1.1-Tr2-RBDv1; RBDv2 – plasmid pTM-RBDv2, MTX concentration in culture medium.

RBDv1 2 uM	RBDv2 2 uM

VCD total, *1000 cells/mL

%viability

| Days in culture | p1.1-Tr2-RBDmyc |
| --- | --- |
| 0 | 293 |
| 1 | 325 |
| 2 | 537 |
| 3 | 862 |
| 4 | 1795 |
| 5 | 2020 |
| 6 | 2305 |
| 7 | 2315 |

|  | % via |
| --- | --- |
| 0 | 82 |
| 1 | 79 |
| 2 | 82 |
| 3 | 80 |
| 4 | 89 |
| 5 | 87 |
| 6 | 76 |
| 7 | 70 |

VCD total, *1000 cells/mL

%viability

| Days in culture | pTM-RBDv2 |
| --- | --- |
| 0 | 493 |
| 1 | 903 |
| 2 | 1736 |
| 3 | 2458 |
| 4 | 3433 |
| 5 | 4370 |

|  | % via |
| --- | --- |
| 0 | 89 |
| 1 | 92 |
| 2 | 93 |
| 3 | 94 |
| 4 | 94 |
| 5 | 90 |

Supporting Figure S2. Cell growth curve for the extended batch cultivation of RBDv1 and RBDv2 – producing cell populations, 2 uM MTX selection pressure. VCD – viable cell density, concentration of live cells – red squares, viability – blue diamonds.

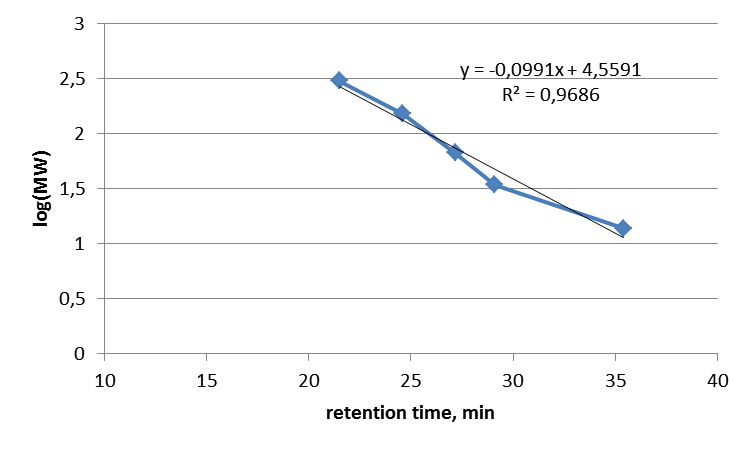

Supporting Figure S3. Size exclusion chromatography trace of molecular mass calibrators and molecular mass calibration curve.

| RBDv1 intact  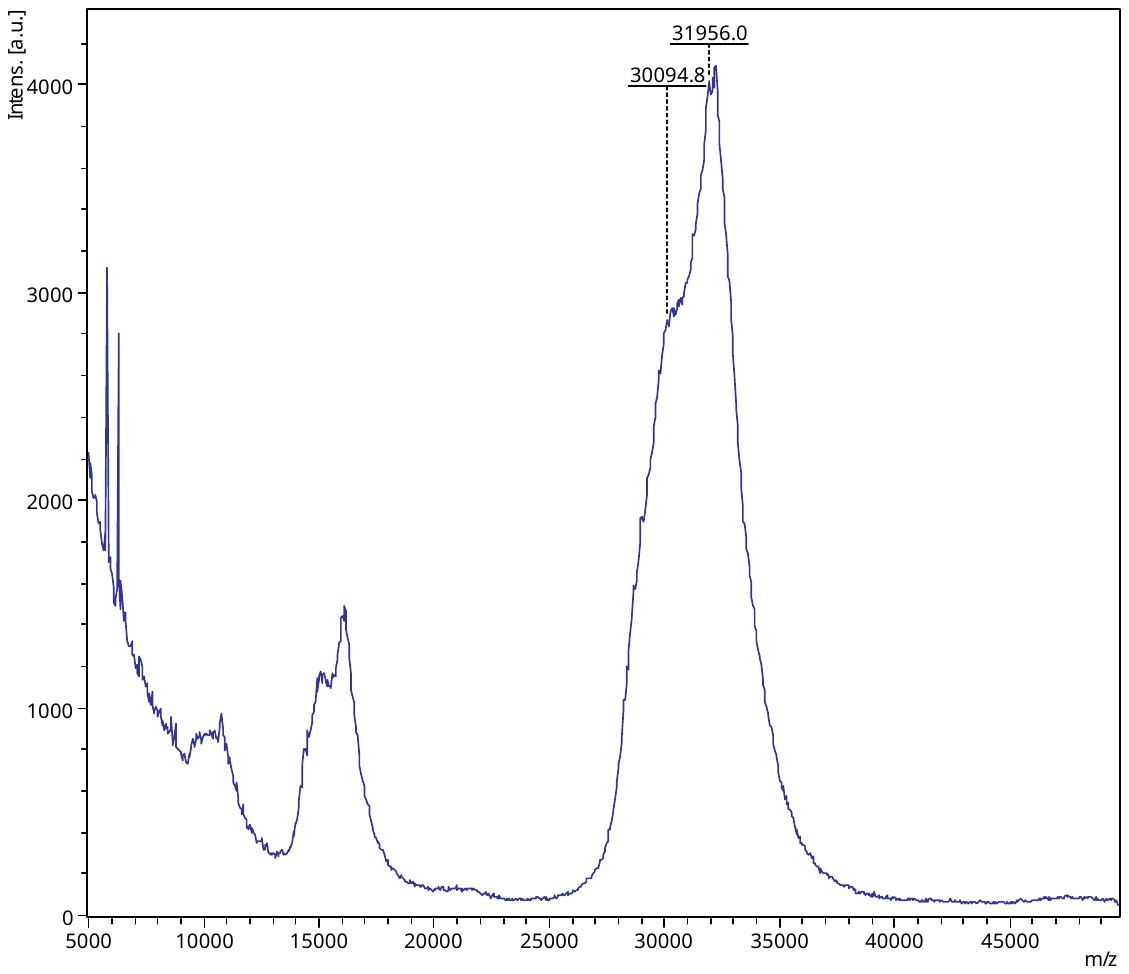 | RBDv2 intact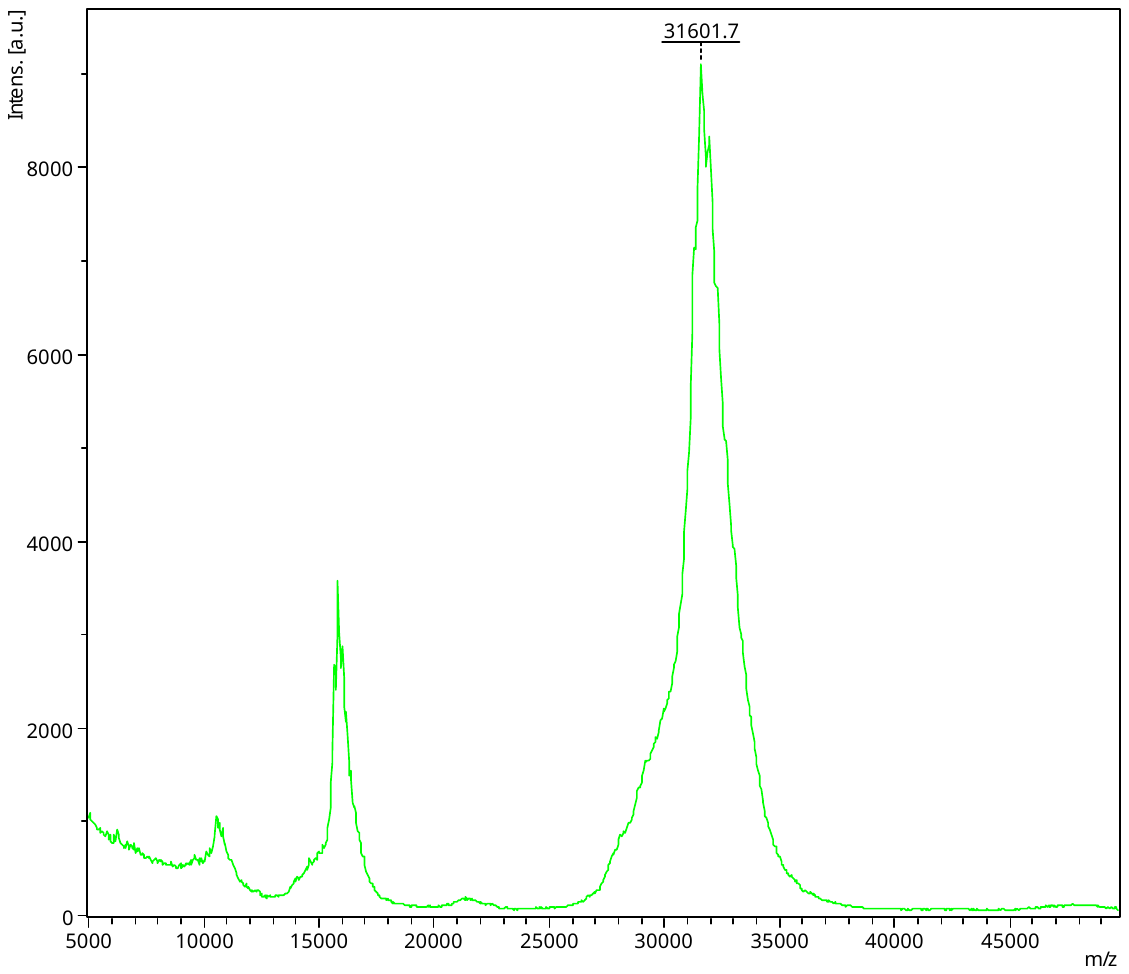 |
| --- | --- |
| RBDv1 deglycosylated  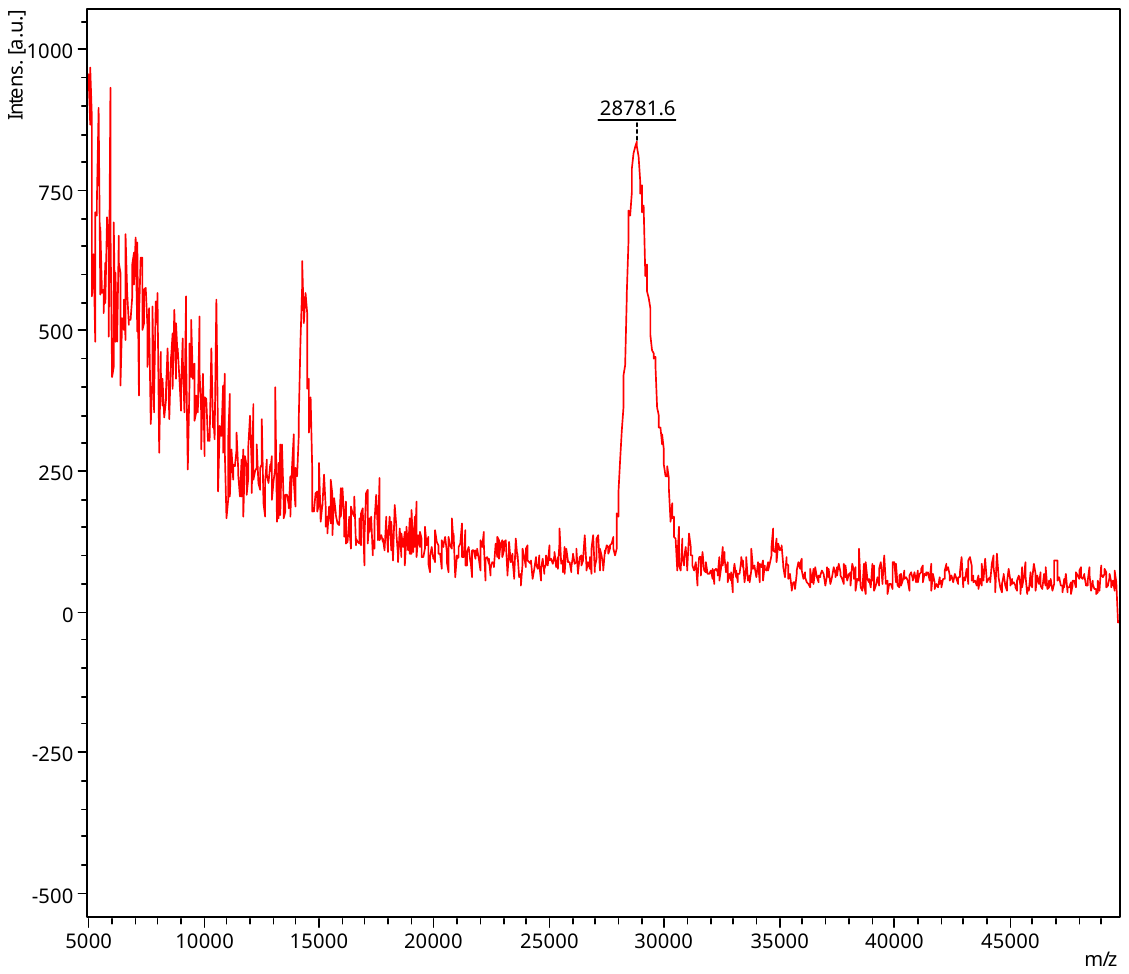 | RBDv2 deglycosylated  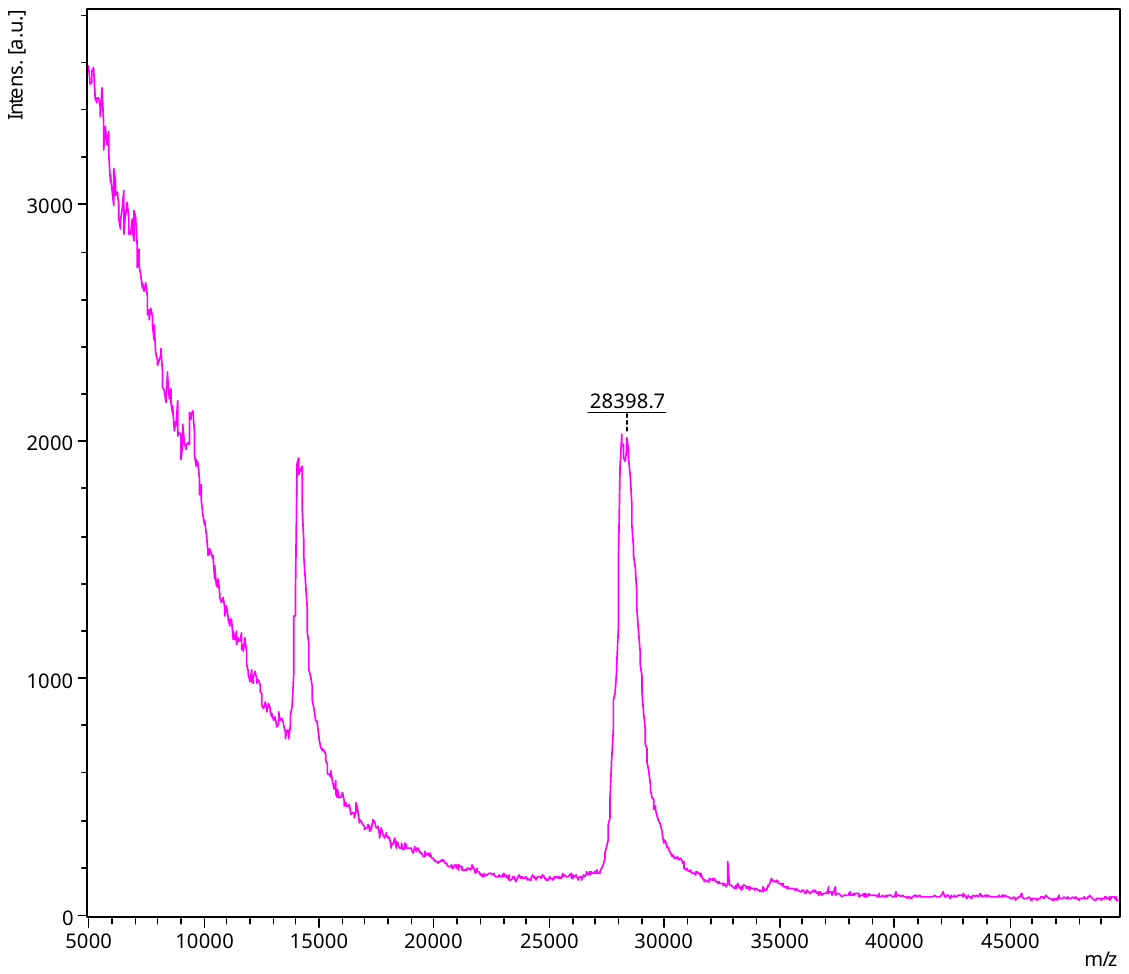 |

Supporting Figure S4. MALDI-TOF spectra traces of intact proteins in glycosylated and deglycosylated forms. Deglycosylated proteins with reduced Cys residues.

RBDv1 intact

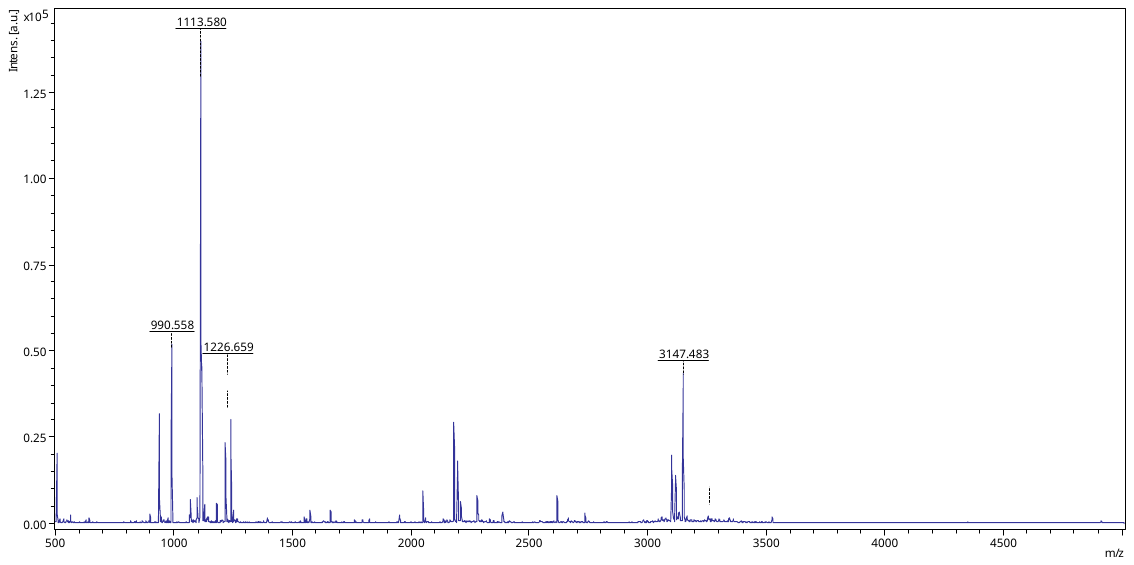

RBDv1 deglycosylated

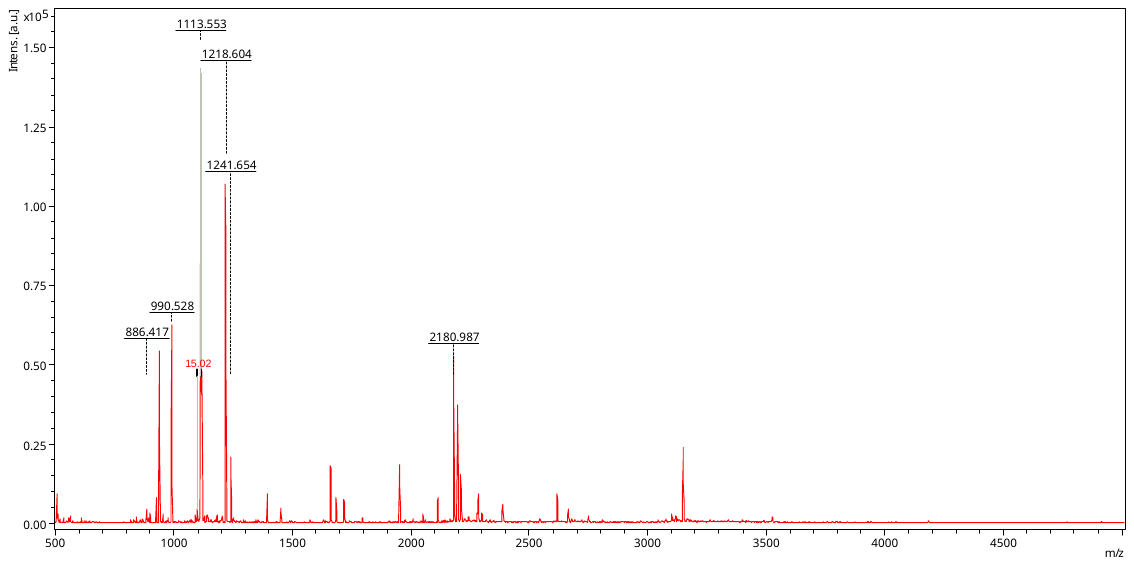

Supporting Figure S5. MALDI-TOF spectra traces of tryptic peptides mxtures from intact and deglycosylated RBDv1. In-gel digestion, reducing conditions, Cys residues not blocked.

RBDv2 intact

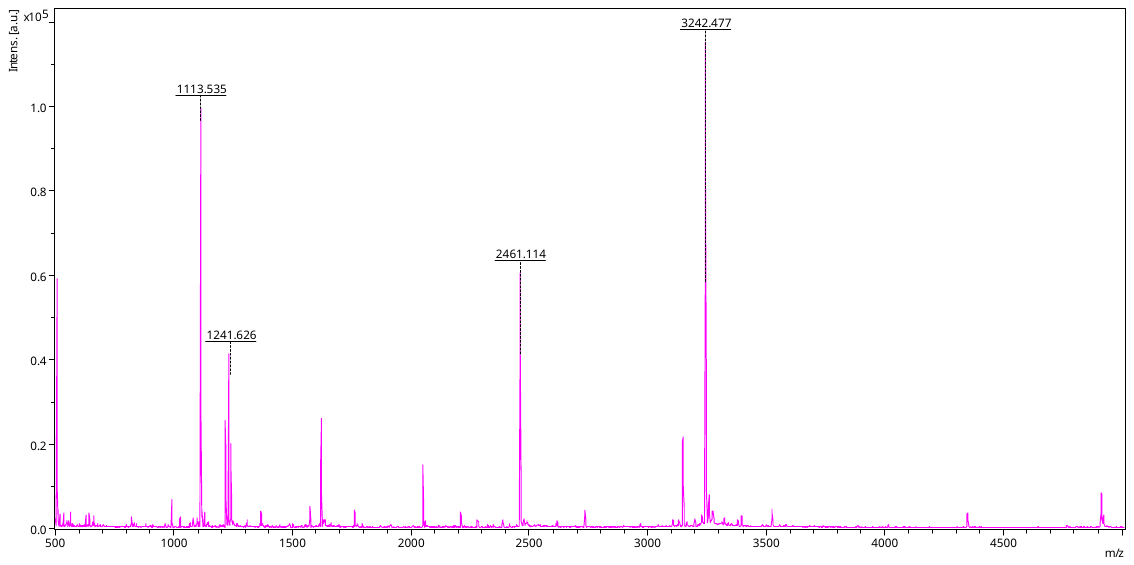

RBDv2 deglycosylated

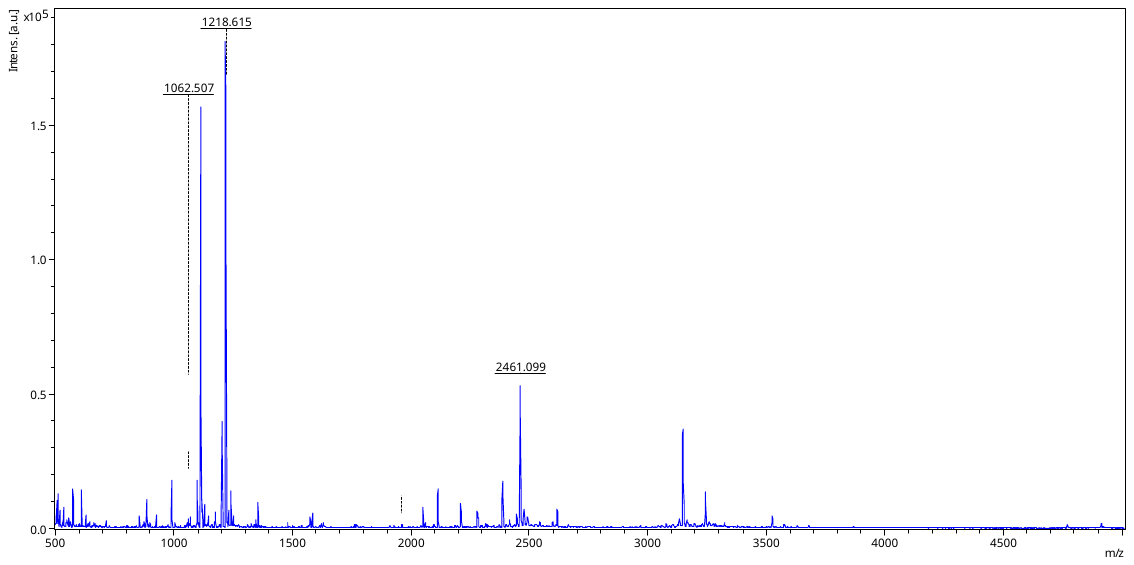

Supporting Figure S 6. MALDI-TOF spectra traces of tryptic peptides mxtures from intact and deglycosylated RBDv2. In-gel digestion, reducing conditions, Cys residues not blocked.

Supporting Table S 1. Peptides mass list of the RBDv2 intact protein, in-gel digestion, reduced protein. Identified peptides are marked by green, peptides not presented on mass list of the RBDv1 – PNGase F spectrum are marked by yellow, non-unique peptides – by gray.

| **m/z** | **S/N** | **QualityFac.** | **Res.** | **Intens.** | **Area** | **start** | **end** |
| --- | --- | --- | --- | --- | --- | --- | --- |
| 789.46 | 5.5 |  | 13409 | 566.00 | 45 | 225 | 231 |
| 818.405 | 6.8 | 606 | 9708 | 673.74 | 97 |  |  |
| 867.41 | 7.6 | 566 | 12320 | 769.04 | 95 |  |  |
| 892.412 | 4.9 |  | 11319 | 517.00 | 47 |  |  |
| 899.502 | 29.5 | 12480 | 12356 | 2954.59 | 402 | 105 | 113 |
| 938.45 | 309.3 | 21576 | 10969 | 30564.47 | 4926 | 234 | 240 |
| 990.558 | 502.0 | 37761 | 9330 | 49870.94 | 10305 | 155 | 162 |
| 1067.56 | 22.8 | 1119 | 13119 | 2334.92 | 379 |  |  |
| 1113.58 | 1283.9 | 43779 | 9557 | 128601.87 | 31309 | 43 | 51 |
| 1118.661 | 428.2 | 44194 | 9959 | 43036.37 | 10164 | 154 | 162 |
| 1129.57 | 47.5 | 13732 | 12729 | 4850.40 | 895 |  |  |
| 1180.601 | 59.9 | 10168 | 15872 | 6139.85 | 1019 | 232 | 240 |
| 1218.618 | 220.9 | 22899 | 11763 | 22903.52 | 5246 | 141 | 150 |
| 1241.669 | 279.0 | 43133 | 13659 | 28900.49 | 5921 | 43 | 52 |
| 1393.735 | 14.3 | 3203 | 14329 | 1553.84 | 372 |  |  |
| 1397.764 | 8.9 | 421 | 16317 | 971.68 | 198 | 43 | 53 |
| 1534.907 | 6.5 | 736 | 15097 | 742.08 | 193 | 151 | 162 |
| 1630.81 | 5.6 |  | 16913 | 676.00 | 86 |  |  |
| 1660.865 | 31.1 | 32685 | 17198 | 3507.65 | 974 |  |  |
| 1684.816 | 6.0 | 1091 | 16219 | 719.26 | 211 |  |  |
| 1762.955 | 8.2 |  | 22822 | 990.00 | 130 | 141 | 154 |
| 1823.951 | 8.0 | 794 | 20483 | 966.72 | 271 |  |  |
| 1951.967 | 17.2 | 2306 | 9352 | 1966.49 | 1293 |  |  |
| 2051.166 | 74.6 | 40034 | 20307 | 8313.85 | 2777 | 206 | 224 |
| 2061.039 | 11.1 | 2465 | 18443 | 1266.61 | 453 | 83 | 99 |
| 2180.988 | 220.0 | 60288 | 17172 | 23389.83 | 9740 | 241 | 258 |
| 2196.975 | 125.7 | 26123 | 15737 | 13324.36 | 6116 |  |  |
| 2280.063 | 61.6 | 12182 | 20478 | 6402.35 | 2418 | 121 | 140 |
| 2387.128 | 25.0 | 6400 | 21439 | 2580.67 | 991 | 54 | 74 |
| 2415.116 | 3.5 |  | 23677 | 541.00 | 80 |  |  |
| 2543.244 | 3.4 |  | 21168 | 546.00 | 70 | 53 | 74 |
| 2617.283 | 52.8 | 25144 | 19374 | 5175.17 | 2531 | 83 | 104 |
| 2734.451 | 18.2 | 3414 | 17118 | 1774.90 | 1017 |  |  |
| 2980.381 | 5.0 |  | 22633 | 782.00 | 109 |  |  |
| 2996.345 | 3.2 |  | 20413 | 533.00 | 57 |  |  |
| 3042.461 | 4.5 |  | 27493 | 748.00 | 98 |  |  |
| 3100.412 | 151.6 | 11270 | 19950 | 11852.49 | 7454 | 234 | 258 |
| 3116.406 | 94.6 | 8561 | 20763 | 7365.91 | 4444 |  |  |
| 3147.482 | 329.9 | 9284 | 18105 | 24881.42 | 17831 | 114 | 140 |
| 3252.445 | 8.8 |  | 21946 | 1246.00 | 252 |  |  |
| 3342.483 | 3.9 |  | 16897 | 602.00 | 117 | 232 | 258 |
| 3358.538 | 3.1 |  | 12411 | 492.00 | 104 |  |  |
| 3523.664 | 12.7 | 2345 | 13613 | 819.53 | 837 | 75 | 104 |
| 4909.984 | 1.5 |  | 8226 | 121.00 | 29 | 163 | 205 |

Supporting Table S 2. Peptides mass list of the RBDv2 intact protein, in-gel digestion, reduced protein. Identified peptides are marked by green, peptides not presented on mass list of the RBDv2 – PNGase F spectrum are marked by yellow, non-unique peptides – by gray.

| **m/z** | **S/N** | **QualityFac.** | **Res.** | **Intens.** | **Area** | **start** | **end** |
| --- | --- | --- | --- | --- | --- | --- | --- |
| 899.478 | 2.6 |  | 6125 | 664.00 | 95 | 113 | 121 |
| 963.381 | 4.7 |  | 9819 | 1067.00 | 143 |  |  |
| 978.361 | 4.1 |  | 9270 | 934.00 | 119 |  |  |
| 990.513 | 33.6 | 8486 | 8337 | 6310.53 | 1445 | 163 | 170 |
| 1067.492 | 4.9 |  | 10717 | 1085.00 | 114 |  |  |
| 1113.535 | 529.0 | 28832 | 7693 | 95865.50 | 29092 | 51 | 59 |
| 1118.602 | 20.8 | 2675 | 8100 | 3890.41 | 1075 | 162 | 170 |
| 1129.521 | 19.9 | 3992 | 9340 | 3728.18 | 916 |  |  |
| 1241.626 | 102.7 | 11234 | 9515 | 18939.16 | 5485 | 51 | 60 |
| 1363.647 | 5.2 |  | 17036 | 1102.00 | 122 |  |  |
| 1374.596 | 5.7 |  | 15618 | 1196.00 | 168 |  |  |
| 1397.716 | 4.2 |  | 17551 | 897.00 | 109 | 51 | 61 |
| 1488.668 | 5.1 |  | 14440 | 1063.00 | 153 |  |  |
| 1500.714 | 5.0 |  | 14322 | 1043.00 | 129 |  |  |
| 1762.927 | 20.4 | 6608 | 14345 | 3678.71 | 1350 | 149 | 162 |
| 1774.826 | 5.6 |  | 13785 | 1136.00 | 158 |  |  |
| 2051.137 | 77.5 | 36052 | 14050 | 12485.62 | 5921 | 214 | 232 |
| 2061.012 | 6.8 | 1470 | 13183 | 1191.50 | 553 | 91 | 107 |
| 2280.049 | 9.8 | 811 | 10349 | 1495.51 | 1035 | 129 | 148 |
| 2387.093 | 6.6 | 653 | 11514 | 997.12 | 624 | 62 | 82 |
| 2461.114 | 302.1 | 34731 | 8960 | 40574.94 | 38644 | 247 | 267 |
| 2617.222 | 7.1 | 1250 | 9519 | 1119.26 | 895 | 91 | 112 |
| 2734.411 | 19.3 | 10587 | 11521 | 2555.76 | 2103 | 149 | 170 |
| 3147.390 | 106.1 | 28148 | 14003 | 11634.52 | 10418 | 122 | 148 |
| 3242.477 | 557.2 | 55130 | 12851 | 57603.56 | 60100 | 240 | 267 |
| 3256.450 | 18.5 |  | 19992 | 3854.00 | 757 |  |  |
| 3322.418 | 6.2 | 323 | 13139 | 950.88 | 647 |  |  |
| 3523.629 | 20.2 | 4288 | 13782 | 1964.01 | 1968 | 83 | 112 |
| 4346.998 | 23.9 | 1812 | 15675 | 1257.69 | 1561 | 122 | 158 |
| 4767.016 | 1.3 |  | 13186 | 261.00 | 55 |  |  |
| 4911.257 | 8.4 |  | 12749 | 1196.00 | 441 | 171 | 213 |
| 4912.199 | 61.5 | 2543 | 12832 | 2178.17 | 4178 |  |  |
| 4921.178 | 21.7 | 2367 | 12720 | 847.22 | 1506 |  |  |
| 5648.234 | 1.7 |  | 8689 | 277.00 | 100 |  |  |
| 5702.585 | 14.9 | 1739 | 11563 | 367.20 | 797 | 1 | 50 |

Supporting Table S 3. Peptides mass list of the RBDv1 protein, treated by PNGase F, in-gel digestion, reduced protein. Peptides not presented on mass list of the intact RBDv1 spectrum are marked by yellow, non-unique peptides – by gray.

| **m/z** | **S/N** | **QualityFac.** | **Res.** | **Intens.** | **Area** |
| --- | --- | --- | --- | --- | --- |
| 789.467 | 1.6 |  | 3244 | 290.00 | 51 |
| 818.382 | 7.2 | 839 | 5895 | 1036.35 | 242 |
| 867.391 | 10.5 | 571 | 5488 | 1495.02 | 416 |
| 886.417 | 27.9 | 6515 | 6126 | 3899.23 | 1015 |
| 892.386 | 11.4 | 411 | 6355 | 1627.35 | 414 |
| 899.465 | 21.4 | 10439 | 6055 | 3003.92 | 829 |
| 925.428 | 58.4 | 24711 | 6719 | 8160.62 | 2107 |
| 938.434 | 390.7 | 32914 | 5811 | 54288.60 | 16523 |
| 954.433 | 19.7 | 3608 | 7203 | 2798.60 | 706 |
| 990.528 | 416.0 |  | 6156 | 62708.00 | 11222 |
| 1075.538 | 7.2 | 569 | 5687 | 1060.19 | 383 |
| 1113.553 | 1075.4 | 34034 | 5901 | 151415.21 | 59821 |
| 1118.634 | 343.7 | 62201 | 6798 | 48390.47 | 16743 |
| 1129.542 | 14.0 | 1533 | 6842 | 2078.13 | 685 |
| 1180.578 | 10.4 | 527 | 7416 | 1581.53 | 502 |
| 1218.604 | 808.8 | 26423 | 6173 | 115583.14 | 50538 |
| 1241.654 | 157.5 | 90611 | 9139 | 22602.02 | 7054 |
| 1393.724 | 61.7 | 77169 | 10518 | 9127.78 | 3007 |
| 1660.857 | 124.4 | 178768 | 11897 | 18979.59 | 7769 |
| 1684.815 | 44.5 | 31495 | 10592 | 6931.76 | 3157 |
| 1950.968 | 9.4 |  | 21297 | 1765.00 | 253 |
| 1951.942 | 103.6 | 69554 | 11966 | 16086.97 | 8501 |
| 2051.145 | 16.1 | 5774 | 16140 | 2527.47 | 1036 |
| 2060.987 | 4.5 |  | 14291 | 918.00 | 154 |
| 2113.018 | 47.1 | 24715 | 17306 | 7050.33 | 2871 |
| 2180.987 | 291.6 | 37318 | 12704 | 42094.65 | 23800 |
| 2196.986 | 198.4 | 48968 | 13375 | 28519.14 | 15457 |
| 2280.063 | 15.5 | 323 | 15551 | 2345.83 | 1081 |
| 2387.117 | 32.2 | 12380 | 15354 | 4458.79 | 2329 |
| 2543.232 | 4.7 |  | 18025 | 1052.00 | 164 |
| 2617.258 | 50.5 | 77746 | 17071 | 6436.99 | 3539 |
| 2967.416 | 2.0 |  | 13415 | 495.00 | 107 |
| 3042.419 | 2.6 |  | 13387 | 620.00 | 110 |
| 3100.365 | 4.8 |  | 18057 | 1046.00 | 147 |
| 3116.392 | 4.9 |  | 22398 | 1074.00 | 160 |
| 3147.446 | 122.0 | 13132 | 15122 | 11983.59 | 9930 |
| 3523.688 | 10.1 | 1174 | 13009 | 929.12 | 911 |
| 4768.323 | 2.2 |  | 15244 | 283.00 | 70 |

Supporting Table S 4. Peptides mass list of the RBDv2 protein, treated by PNGase F, in-gel digestion, reduced protein. Peptides not presented on mass list of the intact RBDv2 spectrum are marked by yellow, non-unique peptides – by gray.

| **m/z** | **S/N** | **QualityFac.** | **Res.** | **Intens.** | **Area** | **start** | **end** |
| --- | --- | --- | --- | --- | --- | --- | --- |
| 854.416 | 19.7 | 3841 | 9197 | 4332.51 | 706 |  |  |
| 886.437 | 47.8 | 25156 | 9332 | 10274.98 | 1782 | 122 | 128 |
| 899.492 | 9.2 | 1439 | 8603 | 2038.86 | 373 |  |  |
| 925.447 | 22.4 | 5310 | 9682 | 4762.69 | 832 | 83 | 90 |
| 990.540 | 90.9 | 31132 | 10363 | 18479.68 | 3536 |  |  |
| 1005.481 | 9.1 | 655 | 9362 | 1918.47 | 384 |  |  |
| 1062.507 | 16.4 | 6122 | 12582 | 3313.86 | 562 |  |  |
| 1067.553 | 8.0 | 390 | 11803 | 1665.37 | 291 |  |  |
| 1113.571 | 785.4 | 53155 | 8665 | 152864.25 | 41316 |  |  |
| 1118.643 | 55.2 | 20578 | 10154 | 10835.89 | 2525 |  |  |
| 1129.559 | 43.8 | 8785 | 10610 | 8631.30 | 1909 |  |  |
| 1218.615 | 869.8 | 15627 | 5792 | 167465.23 | 77755 | 149 | 158 |
| 1241.663 | 68.9 | 35758 | 11313 | 13419.21 | 3384 |  |  |
| 1346.697 | 15.3 | 3927 | 12727 | 3022.57 | 741 |  |  |
| 1480.757 | 9.1 | 1717 | 11610 | 1797.24 | 558 |  |  |
| 1583.779 | 10.4 |  | 13357 | 2173.00 | 313 |  |  |
| 1584.770 | 28.5 |  | 13645 | 5788.00 | 741 |  |  |
| 1630.772 | 9.5 | 320 | 12622 | 1874.12 | 643 |  |  |
| 1762.910 | 7.1 |  | 13082 | 1535.00 | 248 |  |  |
| 1771.834 | 5.8 |  | 13132 | 1269.00 | 172 |  |  |
| 1774.829 | 4.3 |  | 17515 | 965.00 | 124 |  |  |
| 1911.895 | 4.7 |  | 17371 | 1059.00 | 152 |  |  |
| 1928.958 | 3.7 |  | 14804 | 866.00 | 137 |  |  |
| 1961.955 | 3.0 |  | 14912 | 721.00 | 98 |  |  |
| 2051.171 | 37.0 | 23699 | 16999 | 6704.18 | 2682 |  |  |
| 2061.033 | 9.1 | 2078 | 15560 | 1746.77 | 713 |  |  |
| 2113.038 | 74.4 | 41652 | 16787 | 13005.41 | 5530 |  |  |
| 2280.052 | 28.7 | 3384 | 12416 | 4837.80 | 2927 |  |  |
| 2316.086 | 7.8 | 1911 | 13902 | 1420.38 | 709 |  |  |
| 2324.034 | 5.9 |  | 18236 | 1459.00 | 217 | 108 | 128 |
| 2387.106 | 78.1 | 29412 | 12612 | 12310.13 | 7919 |  |  |
| 2461.099 | 224.5 | 14892 | 9995 | 34279.57 | 29036 |  |  |
| 2475.094 | 14.6 |  | 17174 | 3505.00 | 606 |  |  |
| 2617.271 | 33.1 | 39396 | 12399 | 5029.45 | 3677 |  |  |
| 3147.451 | 183.9 | 13025 | 12222 | 20221.20 | 20830 |  |  |
| 3163.420 | 12.2 | 694 | 9661 | 1539.74 | 1717 |  |  |
| 3242.460 | 64.2 | 3025 | 11136 | 6756.68 | 7868 |  |  |
| 3322.435 | 6.7 | 897 | 9415 | 884.06 | 955 |  |  |
| 3523.697 | 23.2 | 4812 | 11587 | 2086.39 | 2533 |  |  |
| 3574.777 | 4.1 |  | 8868 | 849.00 | 261 |  |  |
| 3864.890 | 1.6 |  | 24973 | 354.00 | 35 |  |  |
| 4766.218 | 1.6 |  | 18849 | 222.00 | 34 |  |  |
| 4767.099 | 2.7 |  | 11778 | 317.00 | 105 |  |  |
| 4911.265 | 3.8 |  | 8397 | 421.00 | 192 |  |  |
| 4912.246 | 10.9 |  | 15290 | 1027.00 | 359 |  |  |
| 4921.094 | 6.4 | 770 | 9109 | 227.40 | 417 |  |  |

Supporting Table S 5. List of identified glycopeptides.

| **experimental mass, [M+H]+** | **glycoform mass** | **Δmass (Dalton)** | **structure** | **theoretical glycopeptide mass** |
| --- | --- | --- | --- | --- |
| **RBDv1** |  |  |  |  |
| **1393.735** | 365.132 | 0.03 | (Hex)_1_ (HexNAc)_1_ | 1393.705 |
| **1684.816** | 656.228 | 0.015 | (Hex)_1_ (HexNAc)_1_ (NeuAc)_1_ | 1684.801 |
| **RBDv2** |  |  |  |  |
| **1480.757** | 365.132 | 0.02 | (Hex)_1_ (HexNAc)_1_ | 1480.737 |
| **1771.834** | 656.228 | 0.001 | (Hex)_1_ (HexNAc)_1_ (NeuAc)_1_ | 1771.833 |
